# Limited diversity of bat-associated RNA viruses in endangered and geographically isolated Christmas Island flying foxes

**DOI:** 10.1101/2025.02.27.640670

**Authors:** Ayda Susana Ortiz-Baez, Karrie Rose, Bethan J. Lang, Jane Hall, Edward C. Holmes

## Abstract

Bats are commonly thought to harbour a high diversity and abundance of RNA viruses, some of which are able to jump species boundaries to emerge in new hosts. However, gaps remain in our understanding of the ecological factors that shape the bat virome and influence the diversity, circulation and persistence of viral infections. Flying foxes (Pteropodidae) are representative of the chiropteran fauna in Australia, holding significant ecological, cultural, and social importance. However, some species have also been linked to the circulation of mammalian pathogens such as Hendra virus and Australian bat lyssavirus. Here, we characterised the virome of the Christmas Island flying fox (*Pteropus melanotus natalis*), an endemic and endangered species only found on the remote Australian territory of Christmas Island. Through metatranscriptomic sequencing of 46 samples, including faeces, blood, urine and tissue lesions, we found that these bats exhibit limited RNA virus diversity dominated by dietary viruses. The paucity of RNA viruses likely results from their small population size (between 1500 and 2600 individuals) and the virtual geographic isolation from other bat and mammalian species, except for pests and humans. However, we identified a novel alphacoronavirus in urine, related to viruses circulating in microbats in mainland Australia, and a picorna-like virus related to picornaviruses found in invertebrates. Although this novel picorna-like virus may be of a dietary origin, these flying foxes predominantly eat nectar, pollen and fruit, and viral RNA was also present in blood, urine and wing lesion samples, such that a broader tropism cannot be excluded. Overall, these data reveal how ecological factors have a profound impact on RNA virome diversity, highlighting risks to bat conservation, and showing that bats are not always persistent reservoirs for zoonotic viruses.

## 1. Introduction

Bats (order Chiroptera) are important reservoirs for zoonotic RNA viruses with both epidemic and pathogenic potential, including those within the families *Coronaviridae, Rhabdoviridae, Filoviridae*, and *Paramyxoviridae* [1–6]. Comparative studies of mammalian viromes also indicate that bats harbour a high diversity of viruses relative to many other mammalian orders, and are commonly reservoir hosts in cross-species virus transmission events that are associated with disease emergence [7–10]. In Australia, for example, flying foxes (*Pteropus* spp., family Pteropodidae) have been associated with the circulation and transmission of Hendra virus, Australian bat lyssavirus, and Menangle virus, all of which can cause severe disease or even death in humans and peridomestic animals [2,11–13]. However, although flying foxes have been associated with pathogenic viruses, it is unclear to what extent these animals develop disease due to viral infection, particularly it has been suggested that unique aspects of the bat immune system allow them to tolerate viruses in the absence of overt disease [14–16]. Indeed, recent research has shown that flying foxes carry diverse mammalian-associated viruses with unknown disease associations and zoonotic potential, including paramyxoviruses, caliciviruses, coronaviruses, and retroviruses [17–19].

Christmas Island is a remote Australian territory located in the Indian Ocean, about 350 km south of Java, Indonesia and 2600 km from the Australian mainland, with an area of approximately 135 km^2^. The island has a tropical climate characterized by high temperatures and high humidity throughout the year, as well as pronounced wet (November to June) and dry (July to October) seasons. The insular territory also encompasses rainforests, mangrove forests, and coastal habitats, that support a diverse array of unique fauna and flora. The Christmas Island flying fox (*Pteropus melanotus natalis*, Thomas, 1887; abbreviated here as CIFF) is the last remaining mammalian species endemic to the island. All other endemic mammalian species, including the Christmas Island Pipistrelle bat (*Pipistrellus murrayi*) have been declared extinct likely due to predation, habitat loss and the presence of invasive species. According to the International Union for Conservation of Nature (IUCN), *P. melanotus* is categorized as a vulnerable species, while *P. m. natalis* has been designated as critically endangered [20,21].

CIFF exists as a small and isolated island population. Current estimates of its population size fluctuate between 1500 and 2600 individuals, reflecting a dramatic decrease of 35–75% since the first records in 1980’s [22,23]. This sits in marked contrast to the large population sizes of flying foxes on the Australian mainland where, for instance, surveys of grey-headed flying foxes (GHFF; *Pteropus poliocephalus*), which is classified as a vulnerable species, have resulted in population size estimates ranging between 622,000-692,000 individuals across their habitat in eastern Australia [24], Notably, CIFF populations exhibit low mobility and do not migrate to the mainland or other islands, such that they are isolated from other bat species and mammals more broadly [23]. CIFF are colonial tree dwellers that typically roost in large colonies during the dry season, with peak mating activity occurring between June and August [23,25]. Habitat disturbance is attributed to mining-related activities and urbanization, as well as the introduction of alien species, that exert significant pressure on the population numbers of CIFF, increasing their susceptibility to extinction [26]. CIFF primarily fulfill their nutritional requirements by feeding on fruit and nectar. However, they also consume pollen and foliage, foraging from over 50 plant species and playing a crucial role in pollination and seed dispersal [27].

Despite their endangered status, little is known about the viruses associated with the CIFF [28]. While serological data indicates past natural exposure to viruses such as pararubulaviruses and betacoronaviruses in CIFF [28,29], the lack of molecular evidence makes it challenging to determine the virome composition and the ecology of infection dynamics in this isolated bat population. Herein, we investigated the circulating virus diversity in CIFF by analysing virome composition in a range of tissues, including blood, oropharyngeal swabs and skin lesions, as well as faecal and urine samples, using total RNA sequencing (i.e., metatranscriptomics). We provide evidence that virus diversity in CIFF is very limited, which likely reflects the small population size and isolation of this species that hinders long-term virus maintenance.

## 2. Material & methods

### 2.1 Ethical approval and scientific licenses

The collection of bat samples was approved by the Taronga Conservation Society Australia, Animal Ethics Committee (#3b0423). A permit for access to biological resources from Commonwealth areas was granted by the Australian Government Department of Climate Change, Energy, the Environment and Water (AU-COM2023-583 and PA2023_00070). Samples were imported under permits granted by the Australian Government Department of Agriculture, Fisheries and Forestry (Permit numbers 6761344 & 9871046).

### 2.2 Sample collection

A total of 56 critically endangered Christmas Island flying foxes were collected in August 2023 (i.e., during the dry season) from various locations across Christmas Island (10° 26’ 51” S, 105° 41’ 25” E), Australia. Bats were captured using an aluminium landing net attached to an extendable pole and anesthetized with oxygen and isoflurane (Avet Health Ltd, Lane Cove, NSW, Australia) delivered with a vaporiser and face mask for examination and sample collection. The following samples were opportunistically collected for analysis: swabs from firm, raised, ulcerated, nodular skin lesions on the skin overlying bones of the wing and hind limb (n = 2; Figure 1), an oropharyngeal swab (n = 1), mites (n = 6), blood (n = 22), urine (n = 3) and faeces (n = 12). All samples were preserved in RNA/DNA shield (Zymo Research) according to the manufacturer’s instructions and stored for up to three weeks at -20 ºC during transport, and then at -80 ºC prior to analyses. Bat mites (family Spinturnicidae) were removed from the flying foxes by examining the patagium of the wings and removing mites with fine-tipped forceps. Taxonomic classification of the mites was performed using the *28S* ribosomal RNA gene [30]. Due to the potential co-occurrence of mites and uncertainty in the classification at the species level, the highest achievable taxonomic resolution was at the family level. Punch biopsies (6 mm diameter) of the skin lesions were also collected into 10% neutral buffered formalin, processed in ethanol, embedded with paraffin, sectioned, stained with hematoxylin and eosin, and mounted with a cover slip for examination by light microscopy.

**Figure 1.**
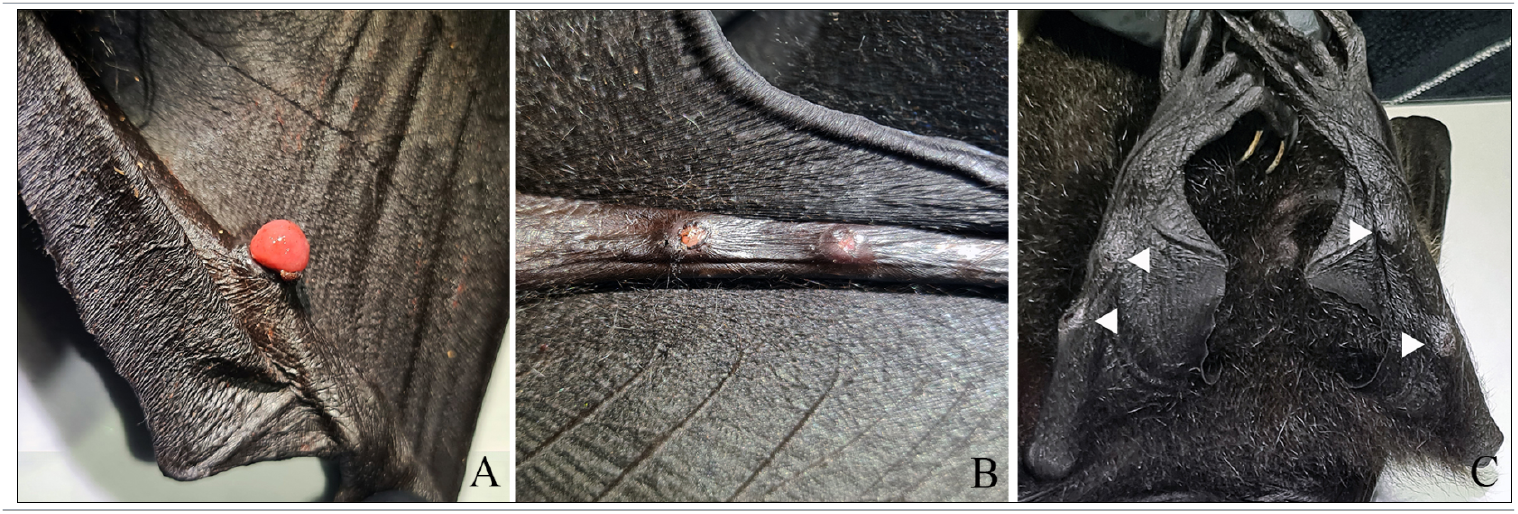
Proliferative and ulcerative lesions observed on the skin of the wing and hind leg of Christmas Island Flying foxes (A-C). White arrow heads illustrate smaller, contracted lesions with central erosion, suggestive of chronicity.

### 2.3 Sample processing and sequencing

Individual bat samples were processed into 43 libraries and homogenised in a lysis buffer using TissueRuptor II (QIAGEN). Briefly, tissue disrupted samples were centrifuged at maximum speed for 15 minutes to remove debris and contaminants from the supernatant. RNA was extracted using the RNeasy Plus kit (QIAGEN, Hilden, Germany) and Quick-RNA Whole Blood kit (Zymo research, USA) according to the manufacturer’s protocol and including negative controls. The total RNA was assessed for its concentration and quality using DeNovix (DeNovix Inc.) and Qubit 3 fluorometer (Thermo Fisher Scientific). Pairedend libraries were constructed using the Illumina Stranded Total RNA with RiboZero Plus kit (Illumina, San Diego, CA, USA) and sequenced on the Illumina NovaSeq X 10B platform..

### 2.4 Sequence data processing and assembly

Quality control of the sequencing data and detection of adapter contaminants was performed using FastQC [31]. Raw sequence reads were pre-processed by trimming low-quality bases and removing adapter sequences using Trimmomatic v.0.39 [32]. The trimmed reads were *de novo* assembled into contigs using Megahit v1.1.3 with default settings [33], and the corresponding open reading frames (ORFs) were predicted on contigs above 900 nt with the getORF program (−minsize 600 -find 0) (EMBOSS software suit). Sequence similarity searches were performed on the resulting contigs and ORFs using DIAMOND BLASTX v.0.9.24 [34] by comparing against the NCBI non-redundant database (NCBI-nr) with an e-value threshold of ≥1×10E-4. To discard false-positives and for further validation of the viral hits, we searched the contigs against the NCBI non-redundant nucleotide database (NCBI-nt) using BLAST with an e-value threshold of ≥1×10E-10 [35]. To assess the distribution of the novel natalis bat picorna-like virus (see Results), we conducted a TBLASTN search against the TSA database (taxid 9397).

### 2.5 Additional targeted microbes

To identify microbes other than viruses, individual libraries were screened *in silico* for the presence of common bacterial and eukaryotic species in bats, including: bacteria – *Bartonella* spp., *Staphylococcus aureus, Salmonella* spp., *Leptospira* spp., *Rickettsia* spp. and *Anaplasma* spp.; fungi – *Pseudogymnoascus destructans, Histoplasma capsulatum, Cryptococcus gatti* and *Paracoccidioides* spp.; and protozoans – *Trypanosoma* spp., *Leishmania* spp., and *Toxoplasma gondii* (Supplementary Table S1). The taxonomic profiling of libraries was performed with Kraken2 v.2.1.1 [36] and MetaPhlAn v4.0 [37].

### 2.6 Domain annotation and phylogenetic analysis

To functionally annotate and detect divergent virus homologs, ORFs and contigs were queried against protein and profile databases using both Interproscan v5.63-95.0 [38] and HMMER v3.3.2 [39]. The A, B and C sequence motifs in the RNA-dependent RNA polymerase (RdRp) were identified using the palmID (https://serratus.io/palmid) and RdRp-scan tools [40,41]. Virus abundance was estimated as the number of transcripts per million (TPM) and reads per million (RPM) with RSEM v1.3.1 [42] and Kallisto v0.46.0 [43]. To account for possible cross-contamination during multiplexing and abundance miscalculation, index-hopping was assumed to occur in cases when read counts were less than 0.1% of the highest count for each virus.

To place the virus sequences obtained within a phylogenetic context, sequences of the RdRp were aligned with reference sequences from the relevant virus taxonomic groups, including the closest hits identified during the BLAST searches, using the E-INS-I algorithm as implemented in MAFFT v7.487 [44]. The best-fit model of amino acid substitution and phylogenetic estimation was then performed using the maximum likelihood (ML) method implemented in IQ-TREE v1.6.12 [45]. Node support was assessed with SH-aLRT and the ultrafast bootstrap (UFBoot, 1000 replicates) [46]. Finally, data visualization was conducted using the R software v4.1.2 and R packages *ape, ggplot2, ggtree, dplyr, RColorBrewer* along with the Inkscape graphic editor software [47,48].

## 3. Results

### 3.1 Virome of the Christmas Island flying fox

Bats are considered important sources of zoonotic viral pathogens due to their unique biology (e.g., physiological traits and social habits), high diversity, and wide geographical distribution [2,5,8,15,49]. However, because virome diversity also reflects ecological factors, such as population size, density and geographic isolation [6,11,50,51], we tested whether the very small population size of the Christmas Island flying fox, as well as its geographic isolation and limited contact with other mammalian species, had impacted the structure of the RNA virome in this species.

Previous studies of *Pteropus* spp. viromes have documented DNA viruses, including those in the families *Herpesviridae, Polyomaviridae, Poxviridae* and *Adenoviridae* [52–57], as well as a wide diversity of RNA viruses, particularly mammalian-associated viruses of the *Paramyxoviridae, Coronaviridae* and *Rhabdoviridae* [28,29,54,58]. Our analysis of the RNA from various tissue and faecal samples from CIFF presented strikingly different results compared with observed in other *Pteropus* spp. Overall, we analysed approximately 9.6 million assembled contigs and identified at least 30 virus families, many within the orders *Picornavirales, Mononegavirales* and *Reovirales* that are associated with invertebrates, plants, fungi, and protozoans based on the known host associations of the closest relatives (Figure 2). The majority of the vertebrate-associated virus sequences detected in CIFF samples were related to various DNA viruses, including different genes from herpesviruses (i.e. Alpha-, Beta- and Gamma herpesviruses) and poxviruses, across all sample types.

**Figure 2.**
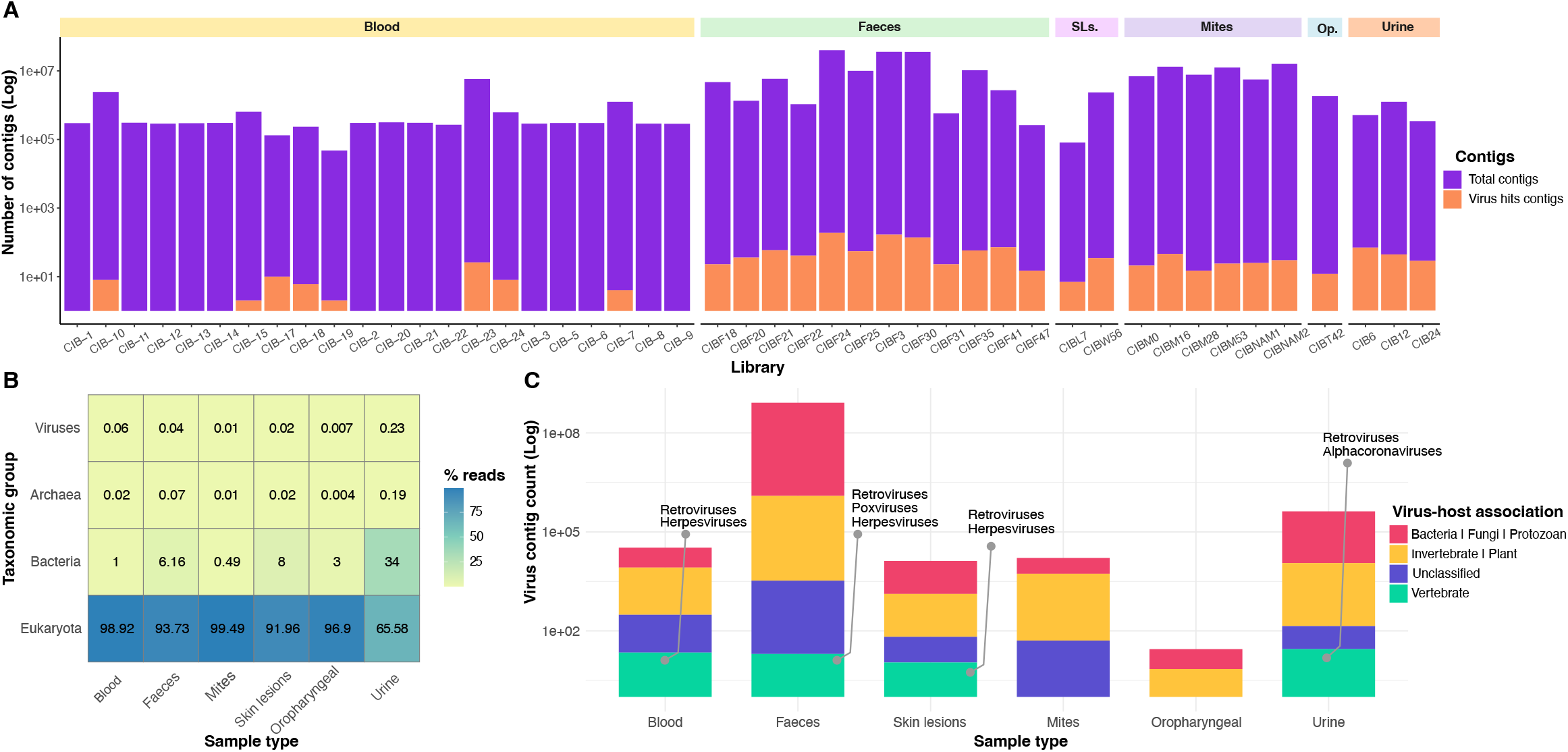
Overview of the RNA virus hits in Christmas Island flying foxes. (A) Distribution of assembled contigs by library, along with the proportion that had a virus hit in the NCBI-nr database. Libraries are grouped by sample type. (B) Taxonomic composition of libraries by sample type, based on the percentage of total reads assigned to domain-level taxa. (C) Viral contig count and composition by host association across sample types. Host assignments are based on virus family classification provided by the ICTV and ViralZone sources. SLs: Skin lesions. Op: Oropharyngeal.

Of most note, the only *bona fide* mammalian-associated RNA virus we identified was a bat alphacoronavirus present in urine (Figure 3; see below). Likely endogenous mammalian retroviral sequences were also detected, including partial sequences related to the murine endogenous retrovirus from the genus *Gammaretrovirus* (∼ 67% amino acid identity) (Figure 2). No RNA viruses were detected in the oropharyngeal swab (CIBT42) and skin lesion (histologically characterised as dermal desmoplasia with epidermal necrosis and ulceration - Figure 1) samples (CIBL7 and CIBW56) aside from retroviruses, plant, and arthropod-associated viruses. Similarly, no viruses likely associated with pathogenic outcomes in bats were identified. Guiyang Paspalum paspaloides tombus-like virus 1 (family *Tombusviridae*; ∼ 50 amino acid identity, e-value= ∼ 1e-137) was detected in both the skin lesion samples and the bat faeces (∼1 – 1360 RPM), although its previous identification in weed samples from China indicates that it is not a mammalian pathogen. Sequences related to leukemia viruses (Gibbon ape and Feline leukemia viruses) and herpesviruses were only found in a single skin lesion sample (CIBL7).

**Figure 3.**
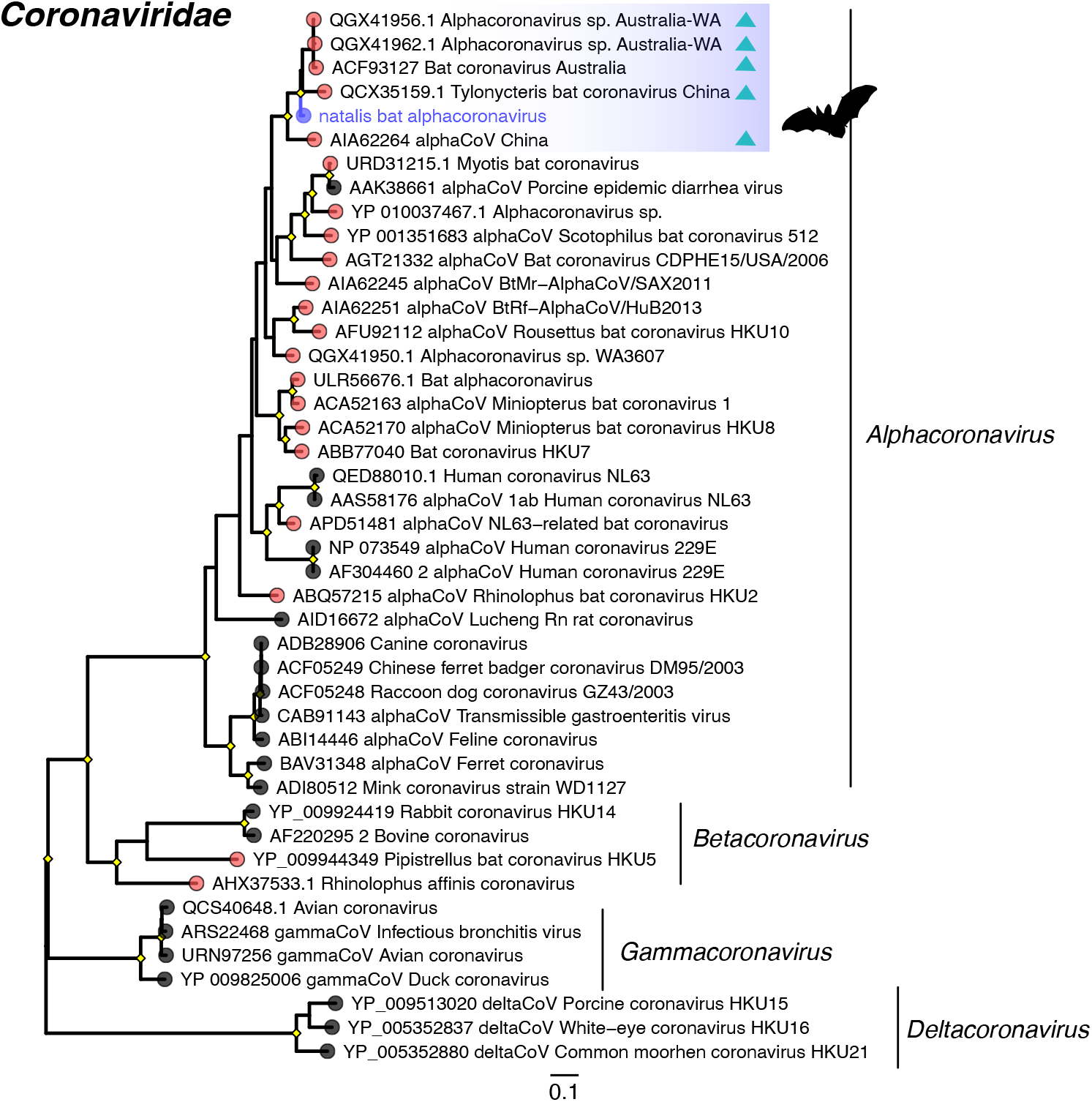
Maximum likelihood phylogenetic tree based on the amino acid sequences of the RdRp protein for RNA viruses within the family *Coronaviridae*. The virus newly identified in this study – natalis bat alphacoronavirus – is shown in purple and grouped within a bat-associated clade as highlighted. Triangles within the bat clade represent microbats (rather than Pteropodidae) from which these viruses were identified. The RdRp tree is midpoint rooted for purposes of clarity. The scale bar indicates the number of substitutions per site. Well supported nodes are denoted with diamond shapes (SH-aLRT >= 80% and UFboot >= 95%). Tips marked in pink across the tree represent viruses identified in bat samples.

As expected, relatively few virus contigs were identified using the NCBI-nr database (Figure 2A). Viral reads within sample types accounted for less than 0.5% of the total reads, with the highest proportion observed in urine and blood samples (Figure 2B). Conversely, the lowest viral contig count was observed in oropharyngeal sample, while the highest was in the faecal samples (Figure 2C).

As noted above, we identified the partial RdRp of an alphacoronavirus in a single urine sample (CIB12). Given its discovery in samples from *P. m. natalis*, this virus was provisionally referred to as “natalis bat alphacoronavirus”. It exhibited 99.2% amino acid identity to an alphacoronavirus found in faecal pellets from Western Australian bats (e-value = 5.66e-159) (Table 1). We utilized this Western Australian sequence as a reference to extend the recovered beyond the RdRp region, which resulted in 85.2% genome coverage (i.e., 23,368 of 27,435 nt), with the number of reads accounting for 0.03% of the entire library (294 RPM). Nonetheless, the genome remained incomplete and contained multiple gaps. Phylogenetic analysis revealed that natalis bat alphacoronavirus clustered within a clade that only includes viruses detected in insectivorous microbats from Australia and China (Figure 3). As such, this represents the first identification of an alphacoronavirus derived from a megabat within this clade.

**Table 1.**
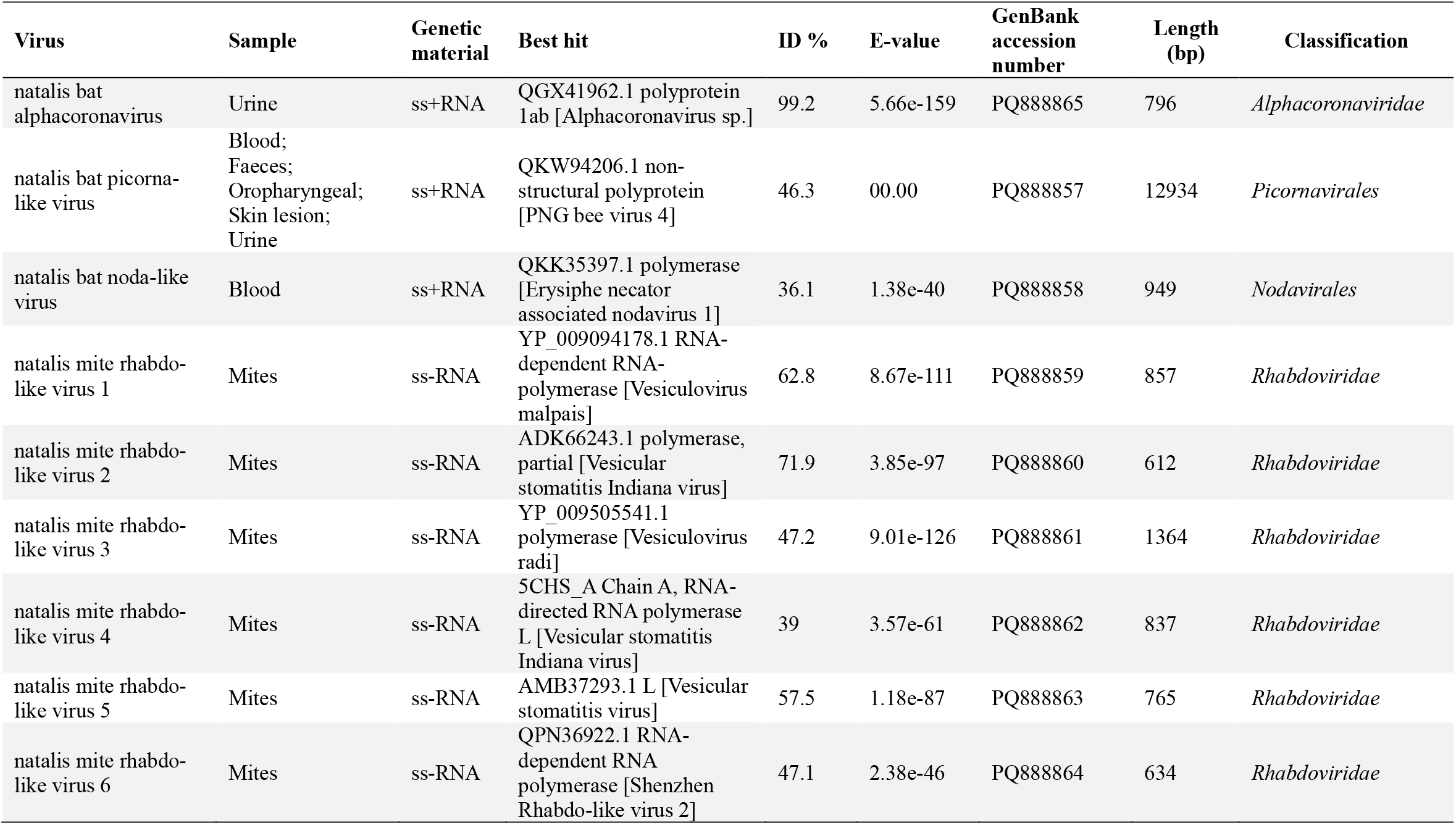
Overview of RNA viruses potentially associated with vertebrate infection, based on top sequence hits and tissue/host distribution.

Also of note was the identification of a novel picorna-like virus (i.e., order *Picornavirales*) in 17 of the 46 libraries, comprising blood, faeces, urine, oropharynx, and wing lesion tissue (Figure 4). No hits were found for this virus in the bat mite libraries. We provisionally named the novel virus as “natalis bat picorna-like virus”. Genomic features of this virus included genes encoding structural proteins for the capsid (VP proteins) and non-structural proteins, such as the glycosyltransferase, RdRp and RNA helicase domains, which are responsible for glycosylation and replication, respectively. Likewise, the RdRp sequence contained the conserved catalytic motifs A (iagDySkFDssl), B (SGsplTsidNSivN) and C (vyGDDnii), as well as the intervening V1 and V2 regions in the canonical order (Palmprint score = 54.3, high-confidence-RdRP). The novel natalis bat picorna-like virus was most closely related to viruses first reported in *Apis mellifera* bees (Figure 4), exhibiting >46% amino acid identity to PNG bee virus 4 (e-value = 4.04E-250), and forming a sister lineage to a clade that includes viruses previously identified in invertebrates, as well as the Wenzhou bat picorna-like virus 8, which was detected in the insectivorous least horseshoe bat (*Rhinolophus pusillus*). Although this phylogenetic position suggests that the natalis bat picorna-like virus is an invertebrate virus, and it was found at relatively high abundance in some of the faecal samples (∼4 – 2900 RPM), it was also present at relatively low abundance level in blood (∼18 – 47 RPM) and at higher abundance in the wing lesion tissue (446 RPM). As such, the possibility that this virus replicates in tissue of the CIFF, a pollen, nectar and fruit-eating species, cannot be fully discounted.

**Figure 4.**
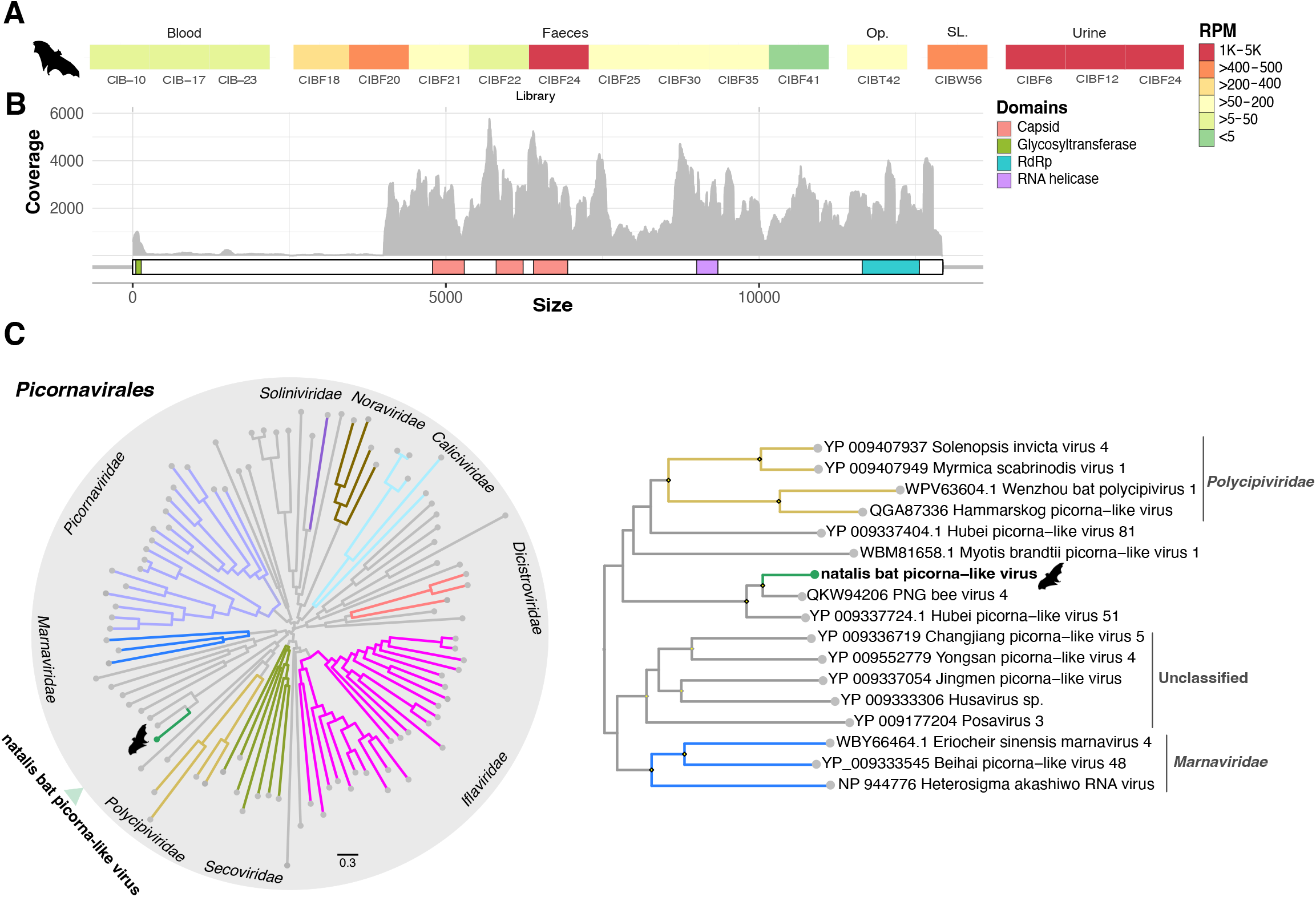
Characterization of the novel natalis bat picorna-like virus. (A) Virus abundance distribution among libraries. Contig quantification is presented in reads per million. (B) Genomic structure and domain annotation. Coverage per position is shown in grey. (C) Phylogenetic tree showing the position of the natalis bat picorna-like virus in the *Picornavirales* (left) with a zoomed-in view of its closest relatives (Right). The RdRp tree is midpoint rooted for purposes of clarity. The scale bar indicates the number of substitutions per site. The branches in grey represent unclassified picornaviruses. Well supported nodes are denoted with diamond shapes (SH-aLRT >= 80% and UFboot >= 95%). SLs: Skin lesions. Op: Oropharyngeal.

In addition, we identified partial polymerase sequences of a novel natalis bat noda-like virus present at low abundance (∼ 11 RPM) in blood sample CIB-17 (Figure 5A, Table 1). This virus was most closely related to fungi-associated viruses based on sequence similarity searches and was distantly placed from vertebrate associated viruses in the phylogenetic tree, suggesting that it is not directly associated with bat infections (Figure 5A).

**Figure 5.**
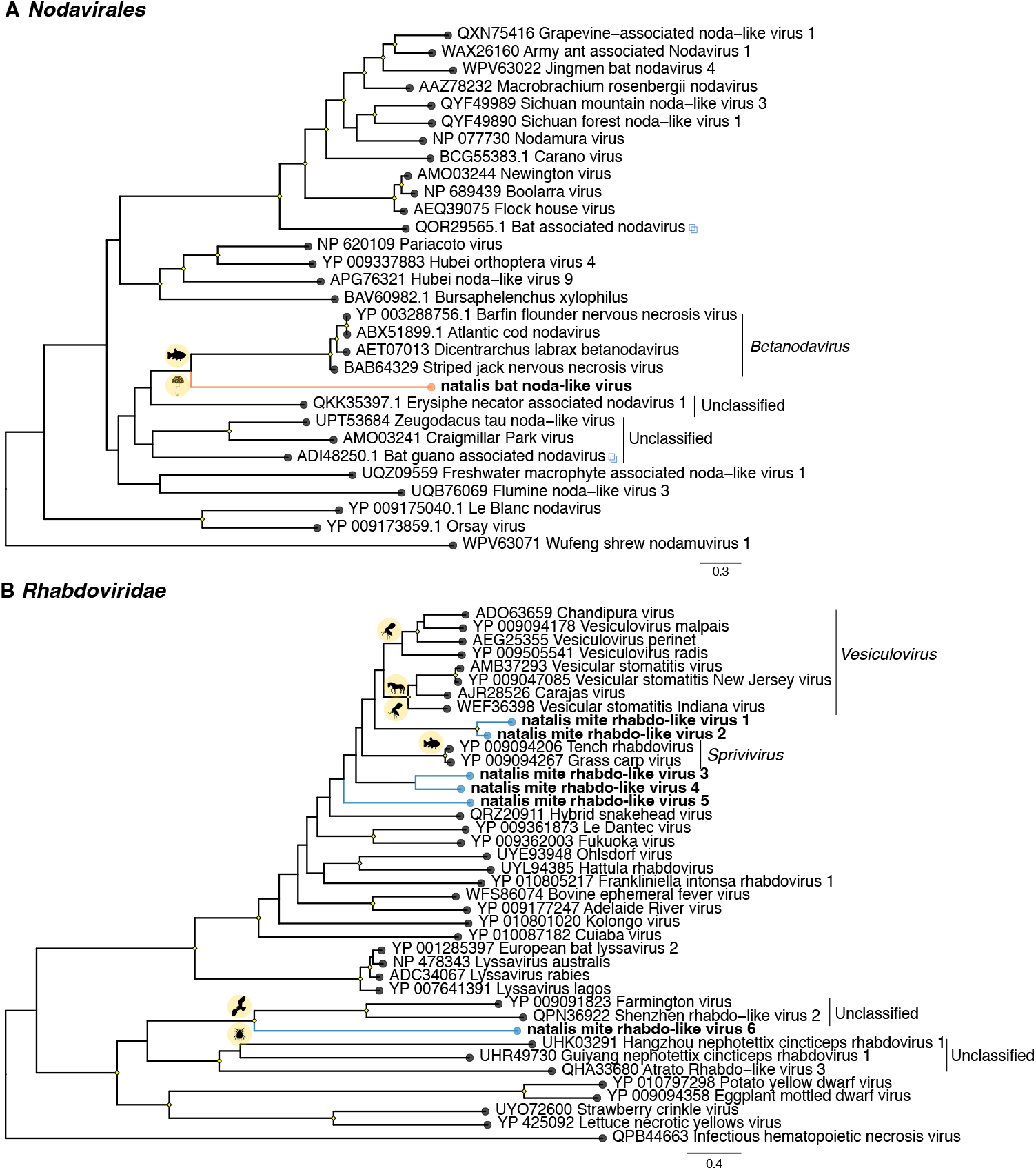
Maximum likelihood phylogenetic trees based on the amino acid sequences of the RdRp protein for RNA viruses within the (A) *Nodavirales* and (B) *Rhabdoviridae*. Viruses newly identified in this study are shown in orange (*Nodavirales*) and blue (*Rhabdoviridae)*. Interlaced squares represent viruses previously identified in metagenomic studies in bat samples. The trees are midpoint rooted for purposes of clarity. The scale bar indicates the number of substitutions per site. Well supported nodes are denoted with diamond shapes (SH-aLRT >= 80% and UFboot >= 95%). Host associations are indicated by silhouettes on the nodes.

### 3.2 Virome of bat mites

The profiling of the bat mite samples revealed mostly arthropod and fungi-associated viruses, and there were no viruses shared between the mites and CIFF tissue samples (Figure 2). Of interest, we detected partial polymerase sequences for various viruses within the *Rhabdoviridae* at almost negligible abundance levels (∼1 RPM), including those related to vesicular disease-associated viruses in the genus *Vesiculovirus*, as well as sequences related to spriviviruses and unclassified viruses (Figure 5B, Table 1).

### 3.3 Bacteria in the Christmas Island flying fox

Finally, despite the ribosomal RNA depletion treatment of the libraries, some common bacterial species were detected, including *Corynebacterium, Streptococcus, Bacteroides, Staphylococcus*, and *Veillonella* species in faecal samples, which are occasionally pathogenic. Similarly, we identified bacteria within the *Staphylococcus aureus*-related complex in the skin lesions (CIBW56). Importantly, however, no definitive causal evidence of pathogenicity was established in this study. Similarly, no targeted fungi or protozoa were identified in CIFF samples.

## 4. Discussion

Research on bats in remote island settings provides an opportunity to explore how ecological interactions shape viral diversity within these environments. In particular, the geographical isolation and endemism of CIFF provides a valuable reference point for studying virus circulation and maintenance in insular systems. Although sampled at a single time point, our data revealed limited virus diversity in CIFF samples that were dominated by diet-derived viruses resulting from the consumption of fruits and pollen, along with the incidental invertebrates present in those plants [59]. In stark contrast to other metagenomic studies of bat populations, only a single mammalian virus was identified, an alphacoronavirus that in turn provides the first molecular (as opposed to serological) evidence of a coronavirus in the endemic CIFF population [29]. The presence of natalis bat alphacoronavirus suggests two possible scenarios: that the virus is endemic to the CIFF population and circulates at low prevalence, or that it was recently introduced from visiting or stranded bats and only circulates transiently. Determining the correct scenario will require more extensive surveys. Notably, natalis bat alphacoronavirus was most closely related to alphacoronaviruses detected in microbats (Microchiroptera) on mainland Australia some 2600 km distant, indicative of cross-species and suborder transmission, although when and where such an event took place is difficult to determine. Since the extinct Christmas Island pipistrelle was the only microbat species inhabiting the island, it is plausible, albeit speculative, that overlapping of habitats or food resources facilitated ecological connectivity and the circulation of alphacoronaviruses in both bat species [60]. Microbat incursion onto Christmas Island from other jurisdictions, however, cannot be excluded.

The discovery of the novel natalis bat picorna-like virus was also of interest. Phylogenetic analysis of this virus clearly places it within a group of invertebrate viruses, suggestive of an invertebrate origin associated with bat diet or through ectoparasites that infest these bats (i.e. hyperparasitism) [61]. However, that the virus was detected in 17/46 samples, including the blood, urine, oropharynx, and skin lesion tissue at relatively moderate abundance (Figure 4), means that replication of the virus in bat cells cannot be entirely excluded and merits further investigation [60]. Indeed, the presence of seemingly invertebrate viruses in mammalian (including human, small mammal and microchiropteran bat) tissues has been observed for viruses within various orders such as the *Picornavirales, Jingchuvirales* and *Hepelivirales* [9,61–64]. It therefore seems possible that these represent the transient replication of an invertebrate virus in mammalian tissue, but without sustained mammalian transmission. Despite the presence of a natalis bat picorna-like virus in one of two skin lesions sampled, its role in pathogenicity remains unclear. The detection of staphylococci in one of the skin lesion samples suggests a possible contribution to lesion development, although likely of an opportunistic nature [65].

In contrast, the discovery of the novel natalis bat noda-like virus at low abundance in a single blood sample, and its phylogenetic distance from fish-associated viruses, suggest an invertebrate origin, likely from dietary sources, and a limited capacity to infect vertebrates [59,61]. Notably, other nodaviruses identified in bats have been detected through metagenomic analysis of guano and clustered with invertebrate-associated viruses, again indicative of a dietary origin [59].

The limited virus diversity found here contrasts with previous findings from serological and molecular studies in CIFF, which provided evidence for the past infection of pararubulaviruses, henipaviruses as well as a betacoronaviruses [28,29], in a similar manner to other *Pteropus* species. Although it has been argued that these viruses might be circulating endemically in the island [29], the small population size of CIFF will result in a low carrying capacity to spread and maintain continuous virus transmission [66], even during the mating season when a higher frequency of contacts is expected. Similarly, seasonality in viral transmission is less compatible with viruses that cause acute infections (e.g. coronaviruses) in small populations due to the transient nature of the infection and the scarcity of susceptible hosts to sustain transmission. Indeed, the herpesviruses and poxviruses present in the CIFF can persist in smaller host populations because of their association with persistent infection and hence an extended transmission period. In addition, the geographic isolation of Christmas Island, and the absence of other mammalian fauna aside from animal pests and humans, mean that seasonal infections could not easily be sourced by other locations or animals. Hence, it is likely that stranded bats might sporadically introduce viral agents into the CIFF population [29,67], leading to waves of infection where the virus enters the population and infects a few individuals. However, the reduced number of hosts would prevent any newly invading viruses from persisting in the population, resulting in the limited presence of mammalian viruses as documented here.

Our study also revealed the presence of rhabdoviruses in *Spinturnicidae* bat mites, but with no evidence of these viruses in the bat samples analysed. This is consistent with previous reports of rhabdoviruses in Spinturnicidae mites, Nycteribiidae bat flies and Trombiculidae chiggers [68,69]. Although bats are known to be an important reservoir for rhabdoviruses, the absence of these viruses in CIFF samples suggests that these rhabdoviruses are of arthropod origin. However, the broad grouping of rhabdo-like virus sequences with those identified in mammals, including vesiculoviruses, which are important zoonotic pathogens in livestock, indicates the potential for these mites to serve as vectors, both transmitting and maintaining rhabdoviruses in bat populations [69–71].

The distinctive characteristics of the CIFF population and the ecological shifts driven by climate and land-use changes mean that the introduction and circulation of potential pathogenic viruses could impact bat conservation on Christmas Island [29]. The relative naivety of the CIFF to a broad spectrum of bat-associated viruses may leave this critically endangered population vulnerable to viral incursions. As such, understanding the role of metapopulation dynamics and critical community size of CIFF in virus transmission will help identify potential threats and address conservation efforts. In this respect, comparative serological profiling of antiviral responses and immunological features between mainland and insular flying foxes, particularly within the framework of longitudinal studies, will enhance our understanding of virus circulation dynamics and the tolerance/resistance mechanisms activated upon virus infection [14].

The association of bats with zoonotic viruses such as Ebola virus and SARS-CoV-2 has bolstered the notion that bats are a consistent and continual virus reservoir [8,15]. In contrast, our study of the Christmas Island flying fox shows how small population sizes and geographic and ecological isolation can have profound impacts on virus diversity. These findings contribute to expand and reframe our understanding of virome diversity and the dynamics of virus infections in bats.

## CRediT authorship contribution statement

A.S.O.B and E.C.H. conceptualized this study. K.R and J.H performed the animal sampling. A.S.O.B and B.J.L. processed the samples. A.S.O.B and E.C.H. analysed the data. A.S.O.B and E.C.H. wrote the original manuscript draft, with contributions from all authors. K.R., J.H, B.J.L and E.C.H. edited and revised the manuscript. E.C.H. supervised the project.

## Data availability

Raw read sequences generated for this project have been deposited at the Sequence Read Archive (SRA) database under Bioproject: PRJNA626677 (BioSample accessions SAMN44448544 – SAMN44448587) and GenBank database (accession numbers: PQ888857 – PQ888865).

## Acknowledgements

Data analysis was conducted using the software available on the University of Sydney HPC Service (Artemis) and the Setonix Supercomputer at the Pawsey Supercomputing Research Centre, Australia. Sampling of Christmas Island flying foxes was made possible through collaboration with Christmas Island National Park personnel, notably Alexia Jankowski, Brendan Tiernana and Kristen Shanygina, and funding from Parks Australia. Taronga Conservation Society Australia is acknowledged for their commitment to wildlife health investigation through long-term support of the Australian Registry of Wildlife Health.

## Supplementary material

**Supplementary Table S1**. List of microbial species included in a custom database for screening libraries using BLASTN.

